# Combining Attention-based Multiple Instance Learning and Gaussian Processes for CT Hemorrhage Detection

**DOI:** 10.1101/2021.07.01.450539

**Authors:** Yunan Wu, Arne Schmidt, Enrique Hernández-Sánchez, Rafael Molina, Aggelos K. Katsaggelos

**Author notes:** https://ivpl.northwestern.edu/, http://decsai.ugr.es/.

## Abstract

Intracranial hemorrhage (ICH) is a life-threatening emergency with high rates of mortality and morbidity. Rapid and accurate detection of ICH is crucial for patients to get a timely treatment. In order to achieve the automatic diagnosis of ICH, most deep learning models rely on huge amounts of slice labels for training. Unfortunately, the manual annotation of CT slices by radiologists is time-consuming and costly. To diagnose ICH, in this work, we propose to use an attention-based multiple instance learning (Att-MIL) approach implemented through the combination of an attention-based convolutional neural network (Att-CNN) and a variational Gaussian process for multiple instance learning (VGP-MIL). Only labels at scan-level are necessary for training. Our method trains the model using scan labels and assigns each slice with an attention weight, which can be used to provide slice-level predictions, and uses the VGPMIL model based on low-dimensional features extracted by the Att-CNN to obtain improved predictions both at slice and scan levels. To analyze the performance of the proposed approach, our model has been trained on 1150 scans from an RSNA dataset and evaluated on 490 scans from an external CQ500 dataset. Our method outperforms other methods using the same scan-level training and is able to achieve comparable or even better results than other methods relying on slicelevel annotations.

## 1 Introduction

Acute intracranial hemorrhage (ICH) has always been a life-threatening event that causes high mortality and morbidity rate [13]. Rapid and early detection of ICH is essential because nearly 30% of the life loss happens in the first 24 hours [18]. In order to prompt the optimal treatment to patients in short time, computer-aided diagnosis (CAD) is being designed to establish a better triaging protocol.

Recently, deep learning (DL) algorithms have been proposed for the diagnosis of ICH. The most direct way is to train models on single slice to detect ICH precisely at slice-level [6, 5]. For instance, Chilamkurthy et al. [6] modified ResNet-18 CNNs to predict ICH of each slice, slice-level probabilities were then combined using a random forest to provide ICH predictions at scan-level.Unfortunately, 3D spatial information is missing when each slice is trained independently. Recurrent neural networks (RNN) were introduced to link the consecutive 2D slices with feature vectors extracted from CNNs so as to enhance sequential connections among slices [3, 12, 14]. Although this approach achieves good performances in ICH detection, its training requires large size hand-labeled datasets at slice-level, whose generation is time-consuming and adds to the burden of radiologists. The use of scan-level annotations greatly reduces this workload. A full scan only needs one single label, which can even be automatically generated by natural language processing (NLP) methods applied to clinical radiologist reports [19]. Therefore, some studies focused on 3D CNNs to predict the existence of ICH at scan-level [19, 9, 17]. However, two major limitations of 3D CNNs are the highly expensive computation and the inability to localize ICH of slice-level which can serve as a instructive guidance for radiologists.

Multiple instance learning (MIL) is a weakly supervised learning method that has been recently applied to DL, especially in the domain of pathology [4]. Here, we treat ICH diagnosis as an MIL problem, where a full scan is defined as a “bag” and each slice in the scan is defined as an “instance”. A scan is classified as ICH if at least one slice in this scan has ICH and is normal if all slices are normal. Few studies use MIL method in ICH detection [15, 16]. For instance, Remedios et al. [15] combine CNNs with MIL to predict ICH at scan-level, but the model was trained with a max-pooling operation so it could only select the most positive instance in a bag.

Variational Gaussian Processes for Multiple Instance Learning (VGPMIL) [7] treat the MIL problem in a probabilistic way. The model has several advantages such as robustness to overfitting (due to the non-parametric modelling provided by Gaussian processes), faster convergence and predictions for instances as well as for bags. One limitation is that the model can not be trained directly on images because of their high feature dimensionality. Therefore we use the attention-based CNN (Att-CNN) as a feature extractor and use the VGPMIL model to make the ICH predictions on slice as well as on scan levels.

To the best of our knowledge, this is the first study that combines Att-CNN for feature extraction with VGPMIL to improve hemorrhage detection using only scan level annotations. We demonstrate that (1) Att-CNN is able to predict accurate slice labels with no need for 2D slice annotations; (2) VGPMIL benefits from Att-CNN and improves the ICH predictions at both slice and scan levels; and (3) our Att-MIL approach outperforms other methods at scan-level and generalizes well to other datasets.

## 2 Methods

### 2.1 Intracranial Hemorrhage Detection as a Multiple Instance Learning Problem

First we mathematically describe the detection of ICH as a problem of multiple instance learning (MIL). We treat CT slices 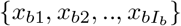 as instances and the full scan as a bag *X_b_*. We assume that all bag labels *Y_b_* ∈ (0, 1)*, b* = 1, 2*, .., B* are available, where *B* describes the total number of CT scans. The true labels of slices 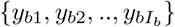 remain unobserved in the multiple instance learning setting. Notice that the number of slices *I_b_* is bag dependent.

When a CT scan contains ICH we know that at least one slice must contain the pattern of hemorrhage while a negative scan contains only negative slices.

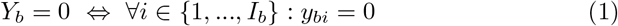

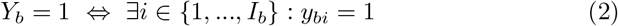

### 2.2 Model Description

The proposed model is defined in Fig.1, training consists of two phases that are executed sequentially. First, the Att-CNN is trained to extract features from the slice images (Phase 1), then the VGPMIL model is trained based on these features to obtain slice and scan level predictions (Phase 2).

**Fig. 1.**
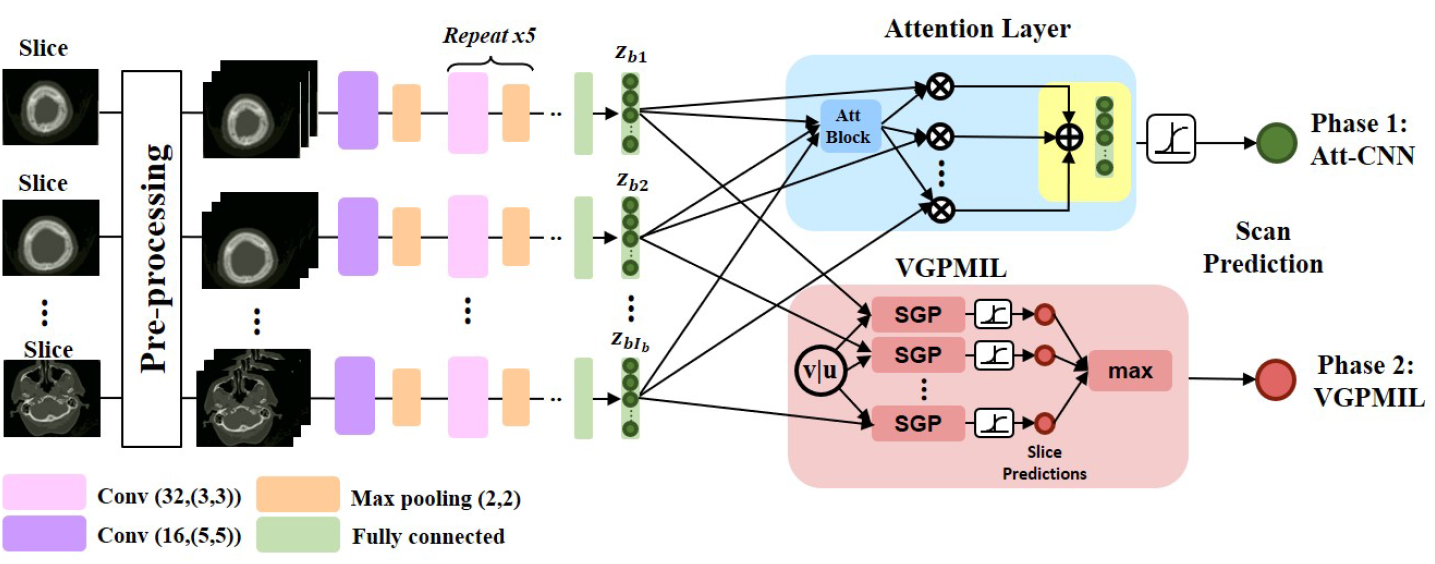
The proposed architecture of Att-MIL, including the Att-CNN in Phase 1 and VGPMIL in Phase 2. Phase 1 updates parameters of Att-CNN at scan-level. Phase 2 uses features extracted from the Att-CNN to train the VGPMIL. The diagnosis are obtained using features of the trained Att-CNN and the VGPMIL for prediction.

**Phase 1** A convolutional neural network *f_cnn_* serves as a feature extractor to obtain a vector of high level features *z_bi_* for each instance *x_bi_* in a bag *b*, so *z_bi_* = *f_cnn_*(*x_bi_*), ∀*i* = 1, 2*, .., I_b_*. The CNN model in Fig.1 is implemented with six convolutional blocks, followed by a flatten layer and a fully connected layer. The convolutional block is used to extract discriminative features from each CT slice, including one convolutional layer and one maxpooling layer. The fully connected layer is used to decrease the size of feature vectors *z_bi_ ∈ R^M×^*^1^, which are fed to the attention layer.

In order to weight each slice differently, we add an attention layer *f_att_* to the CNN (i.e., a two-layered neural networks) [8], where an attention weight *α_bi_* is assigned to each feature vector *z_bi_*. The weights are determined by the model. Specifically, let 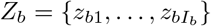 be the set of all feature vectors of bag *b* and 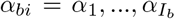, the attention weights for feature vectors *z_bi_*. The weights *α_bi_* add up to 1 and for each bag, there are as many coefficients as instances in the bag. The attention layer *f_att_* is defined as 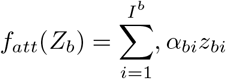, where

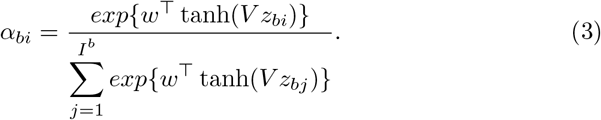

*w* ∈ *R^L×^*^1^ and *V* ∈ *R^L×M^* are trainable parameters, *M* denotes the size of feature vectors and *L* = 50. The non-linearity tanh(·) generates both positive and negative values in the gradient flow. After that, the weighted feature vectors pass through a classifier *g_c_* (i.e., one fully connected layer with sigmoid function) to predict the scan labels:

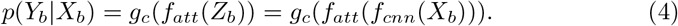

The feature extractor, attention layers and classifier are trained jointly using the common cross-entropy loss until convergence,

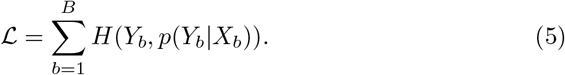

We refer to the whole process as Att-CNN. The weak labels at slice level can be obtained from the attention weights [8, 16]. Specifically, if a scan is predicted as normal, all slices are normal. If a scan is predicted as ICH, the slices with min-max normalized attention weights larger than 0.5 are predicted as ICH. Notice that this is not a proper classifier but a way to obtain one from the attention weights (see [8, 16] for details).

**Phase 2** Once the training at Phase 1 has been completed, we no longer use the attention layer but replace it by a VGPMIL model [7] which has to be trained using *z_bi_* for instance and bag predictions. It is important to note here that in the experimental section we will also analyze the use of *f_att_*(*Z_b_*) as input to VGPMIL (experiment “AL-Aw” in section 3.2).

A Variational Gaussian Process for Multiple Instance Learning is now used to learn to predict slice and CT scan labels (see Phase 2 in Fig 1). Using the trained network, we calculate 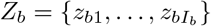, according to *z_bi_* = *f_cnn_*(*x_bi_*). All *Z_b_* are joined together in the matrix *Z*. Then, using the VGPMIL formulation we introduce a set of inducing points *U* = {*u*_1_, …, *u_M_*} and corresponding output *v* which are governed by the distribution 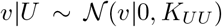, where *K_UU_* is the covariance matrix of the RBF kernel. Using the fully independent training conditional approximation, we utilize the conditional distribution 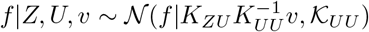, where 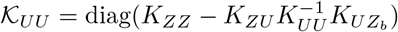. The conditional distribution of *y* = {*y_bi_, b* = 1, …, *B, i* = 1*, …, I_b_*} given the underlying Gaussian process realization *f* is modelled using the product of inde-pendent Bernoulli distributions 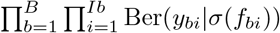, where *σ*(·) denotes the sigmoid function. Finally the observation model for the whole set of bag labels *Y* is given by

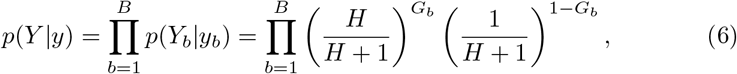

where *G_b_* = *Y_b_* max{*y_bi_, i* = 1*, … , I_b_*} + (1 − *Y_b_*)(1 − max{*y_bi_, i* = 1*, … , I_b_*}) and *H* is a reasonably large value (see the experimental section). Notice that *G_b_* = [*Y_b_* == max{*y_bi_, i* = 1*, …, I_b_*}].

To perform inference we approximate the posterior *p*(*y, f, v*|*Y, Z, U*) by the distribution *q(v)p(f | Z, U, v)q(y)*, with *q(y)* ≔ Π_*b*_ Π_*i*_ *q_bi_(y_bi_)*, which is optimized by solving

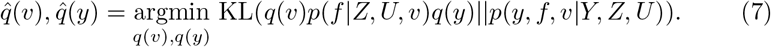

The kernel and inducing location *U* parameters can be optimized by alternating between parameter and distribution optimization. The minimization of the Kullback-Leibler (KL) divergence produces a posterior distribution approximation 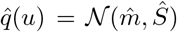, which is used to predict the instance and then the bag labels.

## 3 Experimental Design

### 3.1 Dataset and Preprocessing

A collection of 39650 slices of head CT images acquired from 1150 patients published in the 2019 Radiological Society of North America (RSNA) challenge [1] are included in this study. The number of slices ranges from 24 to 57 for each scan (512×512 pixels). In order to mimic the way radiologists adjust different window widths (W) and centers (C) to read CT images, we apply three windows for each CT slice in a scan with the Hounsfield Units (HU) to enhance the display of brain, blood, and soft tissue, respectively, using [W:80, C:40], [W:200, C:80] and [W:380, C:40]. The three window slices are concatenated into three channels and normalized to [0,1]. The CT scans are split into 1000 (Scan-P:411, Scan-N:589; Slice-P:4976, Slice-N: 29520) for training and validation and the rest 150 (Scan-P:72, Scan-N:78; Slice-P:806, Slice-N: 4448) for testing. Positive(P) represents the case with ICH and Negative(N) represents the normal case. In addition, the models trained on the RSNA dataset are further evaluated on 490 scans (Scan-P:205, Scan-N:285) of an external CQ500 dataset acquired from different institutions in India [6] to test the robustness of the model, each of which has 16 to 128 slices and goes through the same preprocessing steps.

### 3.2 Network Training

The model in Phase 1 is trained with the Adam optimizer [10] with a learning rate of 5 × 10^*−*4^. The batch size is 16 per step. The att-CNN model is trained in two different settings: with no attention layer (nAL), where we use just the unweighted average, and with attention layer (AL). The AL setting is further divided into an experiment where we multiply the extracted features by the attention weight (AL-Aw) and another one with the raw features (AL-nAw) before feeding them into the VGPMIL model. The Att-CNN training process takes an average of about 4.5 hours for 100 epochs with an early stopping operation. We report the mean and standard deviation of 5 independent runs. Both training and testing processes are performed using Tensorflow 2.0 in Python 3.7 on a single NVIDIA GeForce RTX 2070 Super GPU. The VGPMIL in Phase 2 is trained with Numpy 1.19 and runs on the 8 core AMD Ryzen 7 4800HS CPU. All experiments take less than 2 minutes to train the VGPMIL to converge within 200 epochs. The sparse Gaussian processes (SGP) are trained with *H* = 100, 200 inducing points (tested 50, 100, 200, 400) and a radial basis function kernel with a length scale of 1.5 (tested 1.0, 1.5, 2.0, 3.0) and a variance of 0.5 (tested 0.1, 0.5, 1.0, 2.0). The prediction time of the model takes an average of 2.5 seconds to predict a full scan of one patient in both phases.

## 4 Results and Discussions

### 4.1 Attention Layer vs. No Attention Layer

In order to show the capability of the attention layers to learn ICH insightful features, the results in Table 1 provide a comparison of the performances of Att-CNN and VGPMIL with and without attention layers. At scan-level, both Att-CNN and VGPMIL achieve better diagnosis performance with the attention layers. Although the recall is high for results without the attention layers, other metrics are extremely bad. Especially the accuracy score close to 0.5 means that neither the CNN nor the VGPMIL method is able to automatically detect ICH without the attention layers. Similar results are shown at slice-level predictions. Furthermore, we use t-distributed stochastic neighbor embedding (tsne) to reduce the size of the feature vectors at slice-level to two and visualize their distributions to verify our hypothesis. As shown in Fig. 2, it is evident to observe that feature distributions with attention layers (AL) are better separated than those with no attention layer (nAL), which demonstrates the role of attention layers in helping networks learn discriminative features at slice-level.

**Table 1.** Evaluation on the RSNA dataset at slice level and scan level. The results represent the average of 5 independent runs

| Level | Slice-level |  |  |  |  |
| --- | --- | --- | --- | --- | --- |
| Model | Att-CNN+VGPMIL |  |  | Att-CNN |  |
| Metrics | AL-nAw | AL-Aw | nAL | AL | nAL |
| accuracy | **0.938±0.003** | 0.937±0.006 | 0.797±0.005 | 0.923±0.005 | 0.502±0.038 |
| f1 score | **0.786±0.013** | 0.763±0.034 | 0.444±0.030 | 0.773±0.008 | 0.353±0.013 |
| precision | **0.904±0.017** | 0.892±0.018 | 0.382±0.018 | 0.705±0.023 | 0.221±0.010 |
| recall | 0.664±0.024 | 0.668±0.053 | 0.529±0.052 | 0.857±0.015 | **0.884±0.023** |
| Level | Scan-level |  |  |  |  |
| Model | Att-CNN+VGPMIL |  |  | Att-CNN |  |
| Metrics | AL-nAw | AL-Aw | nAL | AL | nAL |
| accuracy | 0.780±0.089 | 0.743±0.176 | 0.603±0.040 | **0.781±0.023** | 0.495±0.009 |
| f1 score | **0.814±0.059** | 0.794±0.104 | 0.707±0.021 | 0.811±0.017 | 0.656±0.004 |
| precision | **0.705±0.099** | 0.705±0.172 | 0.548±0.026 | 0.694±0.023 | 0.487±0.004 |
| recall | 0.975±0.025 | 0.944±0.043 | 0.997±0.006 | 0.975±0.021 | **1.000±0.000** |
| ROC AUC | **0.964±0.006** | 0.951±0.010 | 0.913±0.013 | 0.951±0.011 | 0.768±0.047 |

**Fig. 2.**
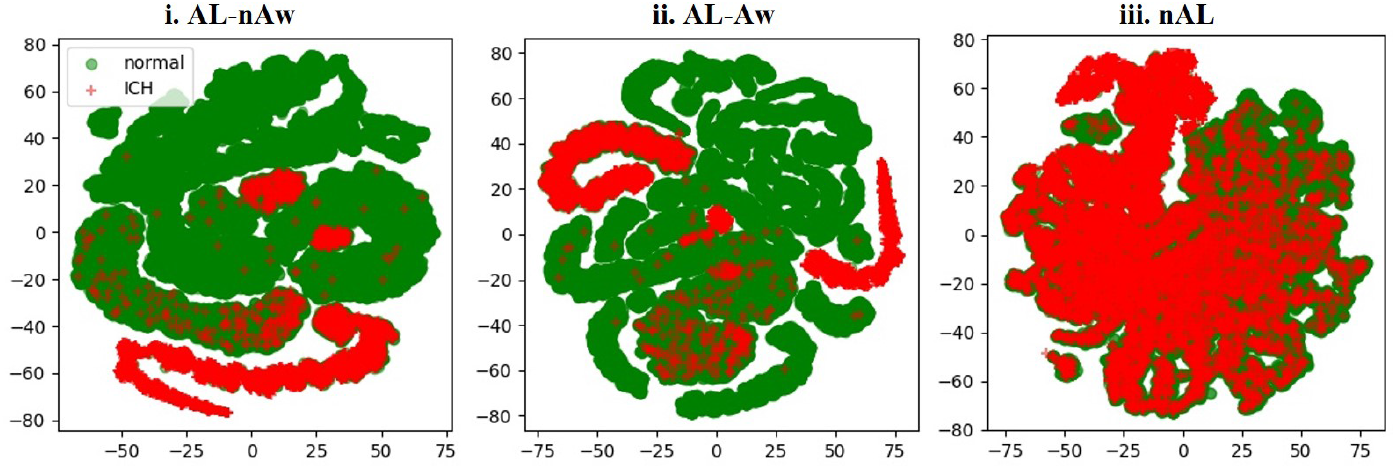
The feature distribution of the RSNA instances in experiment AL-nAw (i), AL-Aw (ii) and nAL (iii). The dimensionality is reduced from 8 to 2 dimensions by t-distributed stochastic neighbor embedding (tsne). We observe that the attention layer (AL) helps the network to learn expressive features for the instances as the classes in AL-nAw and AL-Aw are better seperated than that in nAL.

### 4.2 Attention Weights vs. No Attention Weights

As attention layers are necessary for networks to extract expressive feature vectors, our second hypothesis is that based on those expressive features, VGPMIL is able to achieve good performance even without using attention weights (AL-nAw). In table 1, we compare the results of VGPMIL trained on feature vectors with and without attention weights. At slice-level prediction, VGPMIL without attention weights (AL-nAw) performs slightly better than with attention weights (AL-Aw). At scan-level, both results are equally good, but the overall ROC-AUC of VGPMIL without attention weights (0.964 ± 0.006) is slightly higher than with attention weights (0.951 ± 0.011). The results demonstrate that VGPMIL does not necessarily rely on attention weights to improve ICH predictions because feature separability is present at some level in both cases AL-nAw and AL-Aw, as shown in Fig. 2.

### 4.3 Attention-based CNN vs. VGPMIL

We first do ablation studies with feature dimensions of 8, 32, and 128 for VGP-MIL. The performance is best with 8 features while it gets worse with 32 and 128 features. Next, we do ablations with features extracted from earlier layers (i.e., convolutional layers) where the model performance drops significantly due to their higher feature dimensions. Therefore, VGPMIL chooses the optimal 8-dimension feature size extracted from the fully connected layers of CNN.

Table 1 compares the performance of attention-based CNN and VGPMIL. At scan-level, VGPMIL shows a better AUC score (0.964 ± 0.006) than that of Att-CNN (0.951 ± 0.011). At the slice-level Att-CNN does not provide explicit predictions, but these can be derived from the attention weights as shown in [16, 8]. Therefore, we compare the performance of Att-CNN and VGPMIL at slice-level, where VGPMIL performs better, predicting slice labels at the accuracy of 0.938 ± 0.003 and the precision of 0.904 ± 0.017. This is significant because slice predictions are important for radiologists to localize ICH in a shorter time and the results show that our method is able to infer accurate slice labels even without annotating or training with any slice levels. Table 2 compares the AUC scores of our method with the state of the art training at scan-level. Although these studies use different dataset for their methods, the comparison indicates that our method outperforms them with a relatively smaller dataset.

**Table 2.**
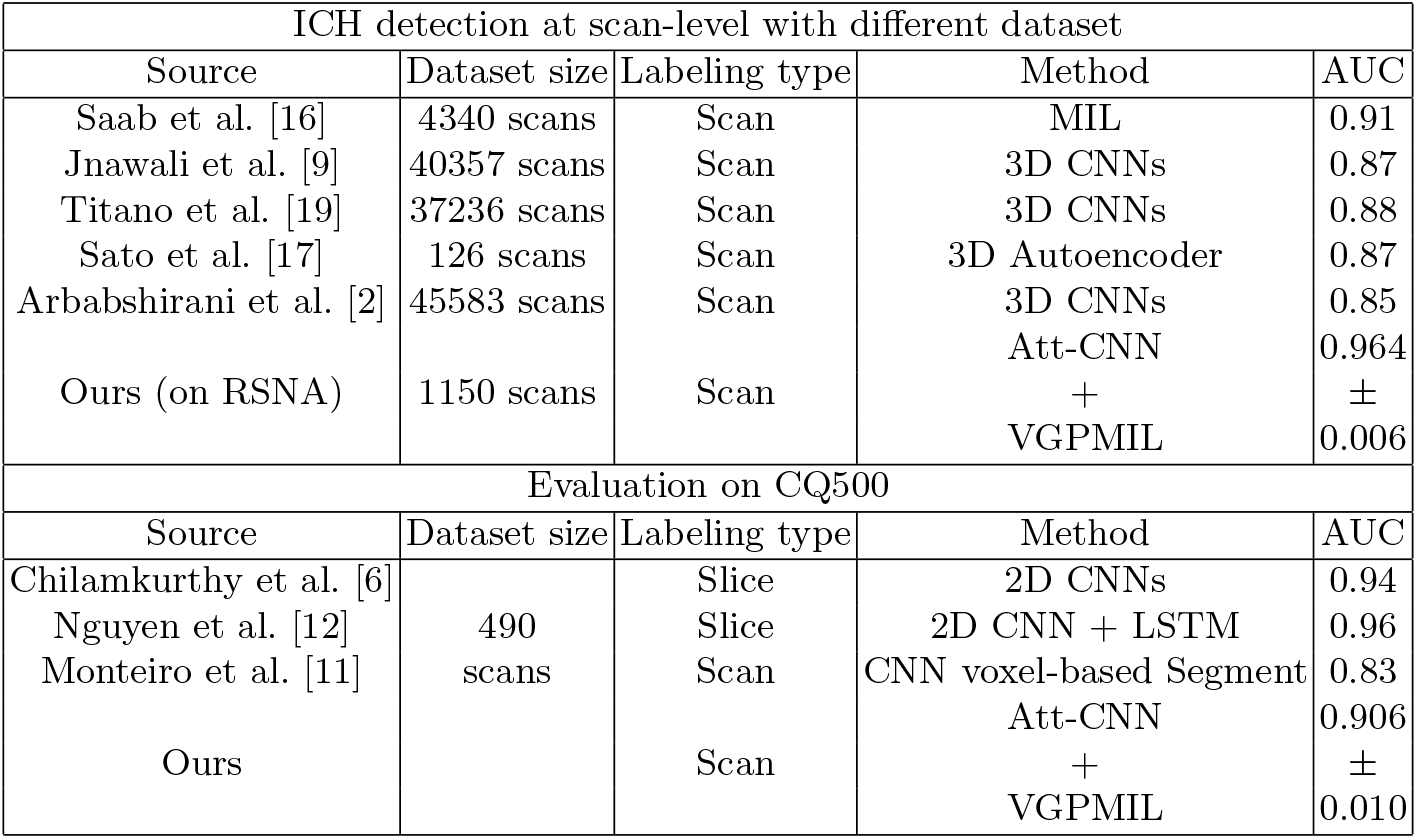
Comparison of different approaches for binary ICH detection. Our results are reported as mean and standard deviation of 5 independent runs.

| ICH detection at scan-level with different dataset |  |  |  |  |
| --- | --- | --- | --- | --- |
| Source | Dataset size | Labeling type | Method | AUC |
| Saab et al. [16] | 4340 scans | Scan | MIL | 0.91 |
| Jnawali et al. [9] | 40357 scans | Scan | 3D CNNs | 0.87 |
| Titano et al. [19] | 37236 scans | Scan | 3D CNNs | 0.88 |
| Sato et al. [17] | 126 scans | Scan | 3D Autoencoder | 0.87 |
| Arbabshirani et al. [2] | 45583 scans | Scan | 3D CNNs | 0.85 |
| Ours (on RSNA) | 1150 scans | Scan | Att-CNN | 0.964 |
|  |  |  | + | ± |
|  |  |  | VGPMIL | 0.006 |
| Evaluation on CQ500 |  |  |  |  |
| Source | Dataset size | Labeling type | Method | AUC |
| Chilamkurthy et al. [6] | 490 scans | Slice | 2D CNNs | 0.94 |
| Nguyen et al. [12] |  | Slice | 2D CNN + LSTM | 0.96 |
| Monteiro et al. [11] |  | Scan | CNN voxel-based Segment | 0.83 |
| Ours |  | Scan | Att-CNN | 0.906 |
|  |  |  | + | ± |
|  |  |  | VGPMIL | 0.010 |

Finally, we evaluate our method on an external testing CQ500 dataset and compare ROC scores with other methods (see Table 2). The labeling type “Scan” means the labels on scan-level in our training dataset, and the type “Slice” means the training labels on slice-level. For comparison, our method outperforms [11] predicting ICH at the same scan-level and is comparable to [6, 12] training on slice labels. Notice that training on slice-level [6,12] is easier as it is fully supervised and involves more labels than scan-level methods. The results on CQ500 dataset further prove the good generalization of our method.

## 5 Conclusions

In this work, we propose an attention-based MIL method that combines attention-based CNN and VGPMIL to predict ICH at scan-level and achieves competitive AUC scores to previous works. Attention layers are important to extract meaningful features that VGPMIL relies on to improve the ICH predictions at both scan and slice levels. Importantly, our method is able to accurately predict slice labels without any slice annotations, greatly reducing the workload in future clinical research. Furthermore, the evaluations on the external dataset prove the good generalization of our method. This study paves the way for a promising weakly supervised learning method that fuses VGPMIL in CNN as a single end-to-end trainable model.

